# Early life stress causes persistent impacts on the microbiome of Atlantic salmon

**DOI:** 10.1101/2020.01.16.908939

**Authors:** Tamsyn M. Uren Webster, Sofia Consuegra, Carlos Garcia de Leaniz

**Affiliations:** Centre for Sustainable Aquatic Research, College of Science, Swansea University, Swansea, SA2 8PP, UK

## Abstract

Intensively farmed fish are commonly stressed, often leading to immune impairment and increased susceptibility to disease. Microbial communities associated with the gut and skin are vital to host immune function, but little is known about how stress affects the fish microbiome, especially during the sensitive early life stages. We compared the effects of two aquaculture-relevant stressors on the gut and skin microbiome of Atlantic salmon fry: an acute cold stress during late embryogenesis, and a chronic environmental stress during the larval stage. Acute cold stress had a lasting effect on the structure of both the gut and the skin microbiome, likely due to disruption of the egg shell microbial communities which seed the initial colonisation of the teleost microbiome upon hatching. In contrast, chronic post hatch stress altered the structure of the gut microbiome, but not that of the skin. Both types of stressors promoted similar Gammaproteobacteria ASVs, particularly within the genera *Acinetobacter* and *Aeromonas* which include several important fish pathogens and, in the gut, reduced the abundance of Lactobacillales. This suggests that there may be common signatures of stress in the salmon microbiome, which may represent useful stress biomarkers in aquaculture.

## Introduction

Microbial communities associated with the gut, skin and other mucosal surfaces have a fundamental influence on host fitness, including nutrient acquisition, metabolism and immune competence (Koskella et al. 2017). In particular, the microbiome plays a critical role in the maturation of the vertebrate adaptive immune system and the stimulation of immune response, and can directly enhance host pathogen defence via colonisation resistance and production of inhibitory compounds (Brestoff & Artis 2013; Kamada et al. 2013). Evidence from mammalian studies suggests that host resilience to stress, disease and immune-related disorders is critically dependent on microbiome diversity and functionality (Rea et al. 2016). However, the host-microbiota relationship is finely balanced and sensitive to disruption by environmental stressors (Foster et al. 2017). Stress influences the mammalian brain-gut-microbiota axis, including neural, immuno, and endocrine signalling pathways, via complex and interacting mechanisms (Foster et al. 2017). For example, host stress response mediated via the hypothalamus-pituitary-adrenal (HPA) axis (i.e. secretion of corticosteroids and catecholamines) can directly influence microbiome diversity, structure and function (Huang et al. 2015), while microbiota and their metabolites are also known to exert modulatory effects throughout the HPA system, altering stress response (de Weerth 2017; Simard et al. 2014). Disruption of the gut-brain axis, including microbiome dysbiosis, has been linked to a number of stress-related conditions, including suppression of immune function, depression and anxiety, inflammatory bowel disease (IBD), obesity and metabolic syndrome (Borre et al. 2014; Foster et al. 2017; Rea et al. 2016).

In intensive aquaculture, fish are often reared at unnaturally high densities and exposed to a range of other handling and social stressors, which can induce adverse health effects such as impaired growth and fitness, altered behaviour and depressed immunity (Conte 2004; Ellison et al. 2018; Iwama et al. 2011; Rodriguez-Barreto et al. 2019). Some evidence indicates that aquaculture related stressors can alter fish microbiome diversity and structure (Boutin et al. 2013; Ringø et al. 1997; Zha et al. 2018). Additionally, environmental stressors such as temperature, toxicant exposure or pH may directly affect the microbiome of aquatic species which are in intimate contact with the external environment (Claus et al. 2016; Sylvain et al. 2016; Zarkasi et al. 2014). This suggests that aquaculture has the potential to disrupt fish microbial communities in the gut, as well as on the skin and other mucosal surfaces, which may have consequences for fish health and disease resistance. However, relatively little is known about the impacts of different types of aquaculture-related stress on the fish microbiome, or how these may influence host health.

The early life stages of teleost fish are particularly sensitive to environmental stressors, reflecting developmental plasticity during critical periods for the assembly of the teleost microbiome, as well as maturation of the nervous and immune system (Giatsis et al. 2015; Uren Webster et al. 2018b). In mammals, stress during early life has a critical influence on gut microbial colonisation and community establishment, with long lasting effects on both the microbiome and health of the host (Foster et al. 2017), however the sensitivity of the fish microbiome to early-life stress is unknown. The teleost intestine is colonised upon hatching from microbes present in the surrounding water and attached to egg-shell fragments, and these early communities are very dynamic and readily influenced by environmental variation (Giatsis et al. 2015; Ingerslev et al. 2014). Although diet becomes a dominant factor shaping further proliferation and differentiation of the gut microbiota (Giatsis et al. 2015; Smith et al. 2015), earlier community dynamics may still have significant lasting effects on future microbial colonisation (Sprockett et al. 2018; Walter & Ley 2011). Here we examined the effect of two aquaculture-relevant stressors on the microbiome of Atlantic salmon (*Salmo salar*) during early development. We hypothesised that microbiome diversity and structure would change depending on the timing and nature of the stressor. Therefore, we characterised the skin and gut microbiome of four month old Atlantic salmon following exposure to an acute cold stress applied during late embryogenesis and a chronic post-hatch environmental stress.

## Methods

### Ethics

All experiments were performed with the approval of the Swansea Animal Welfare and Ethical Review Body (AWERB; approval number IP-1415-6).

### Stress experiments

The stress experiments and husbandry conditions are fully described in Uren Webster et al. (2018b). Briefly, Atlantic salmon eggs from 10 families were maintained in vertical incubators, supplied with flow-through de-chlorinated tap water. Hatched larvae were transferred to shallow troughs (100L x40W x 8D) supplied with artificial substrate to provide support for egg sac reabsorption and shelter for alevins until emergence. Fry were fed with a commercial salmonid feed (Nutraplus, Skretting, UK) of the appropriate grade and quantity recommended by the manufacturer. Water oxygen saturation (>90%), ammonia (<0.02 mg/L), nitrite (<0.01 mg/L), nitrate (<15 mg/L) and pH (7.5 ± 0.2) were maintained within appropriate ranges. Water temperature was gradually increased from 9 °C to 11 °C and photoperiod adjusted from 10L:14D to 14L:10D over the four months of the experiments, reflecting seasonal change.

Eggs were randomly assigned to three experimental groups: control, acute cold stress and chronic environmental stress. Each group was maintained in two replicate egg trays/fry troughs, each containing 500 individuals. The acute stress consisted of a cold shock (five minutes immersion in iced water (0.2 °C), followed by five minutes air exposure (12 °C)), during late embryogenesis (360 degree days; DD). For the chronic stress, hatched larvae were maintained in bare fry troughs lacking the artificial substrate provided to supply support during yolk sac reabsorption and shelter for larvae/fry in the other experimental groups throughout the duration of the experiment (from 475-1532 DD). Mortality, hatching success, and size were recorded throughout the experiment. Neither the acute or chronic stressors altered overall hatching success or survival and, although we initially measured a modest reduction (15%) in the weight of chronically stressed fish, there was no difference in final size (length, weight) or condition index at the end of the four month experimental period (Uren Webster et al. 2018b).

### 16S rRNA sequencing & bioinformatics

At the final sampling point (1532 degree days) fish were euthanised via an overdose of anaesthetic (Phenoxyethanol; 0.5 mg/L), followed by destruction of the brain according to UK Home Office regulations. Skin mucus was collected using Epicentre Catch-All™ Sample Collection Swabs (Cambio, Cambridge, UK), by swabbing each fish along the entire length of the lateral line five times in both directions, on the left-hand side of the body. Sterile dissection of the whole intestine was performed to include both the intestinal contents and epithelial associated microbial communities. Skin swabs, intestine samples and 50 ml water samples from each tank were stored in sterile tubes at −80 °C until DNA extraction.

16S rRNA analysis was performed for a total of 10 individuals (five per replicate tank) from each of the three experimental groups (acute stress, chronic stress, control). DNA extraction from the intestine, skin swabs and water samples was performed using a PowerSoil DNA Isolation Kit (Qiagen, Manchester, UK) according to the manufacturer’s instructions. 16S library preparation using Nextera XT Index kit was performed according the Illumina Metagenomic Sequencing Library Preparation guide, amplifying the V4 hypervariable region of the bacterial 16S gene as fully described in Uren Webster et al. (2018a). The primers 519F (5’-AGCMGCCGCGGTAA-3’) and 785R (5’-TACNVGGGTATCTAATCC-3’) were used for skin and water samples but were associated with excessive non-specific amplification of host DNA from the gut, therefore the primers 341F (5’-CCTACGGGNGGCWGCAG-3’) and 785R (5’-GACTACHVGGGTATCTAATCC-3’) were used for the gut samples instead.

Raw sequence reads have been deposited in the European Nucleotide Archive under study accession number PRJEB32293. Analyses of gut, skin and water samples were performed separately, within Qiime2 (v2019.4, (Bolyen et al. 2018)). Raw sequence reads were initially quality screened and truncated to 280 bp (forward reads) and 240 bp (reverse reads), and the first 8 bp were removed to eliminate potential adaptor contamination. Trimmed reads were then de-noised, merged, subject to chimera screening and removal, and assigned into actual sequence variants (ASVs) using DADA2 (Callahan et al. 2016). Taxonomic classification of ASVs was performed using the Silva reference taxonomy (v132; (Quast et al. 2013)) with a custom trained classifier (Bokulich et al. 2018) specific to the primer pair employed. Host mitochondrial sequences and chloroplast sequences were removed from the dataset, and good reads were subsampled to an equal depth (skin and water-6718; gut-2716) before calculation of alpha and beta diversity metrics (Chao1 richness, Shannon diversity, and Bray-Curtis dissimilarity). One gut sample (control) and one skin sample (chronic stress) were eliminated from the analysis due to very low read numbers.

### Statistical analysis

All statistical analysis was performed in R v3.5.0. We employed linear mixed effect models (using the lme4 package) to examine the effects of stress treatment and fish size on measures of alpha diversity in the skin and gut, including tank identity as a random factor. We included fish length as a covariate of size effects because it had a lower coefficient of variation than fish mass (CV= 8% vs 28.2%). In each case we achieved model simplification by performing backward selection using the *step* and *drop1* functions and selected the model with the lowest AIC value. We then refitted the final model using Restricted Maximum Likelihood, or as a linear model when tank identity (random component) did not improve model fit. We examined the effects of stress treatment on alpha diversity in the tank water using linear models. Structural analysis (microbial beta diversity) was based on community distance matrices calculated using the Bray-Curtis dissimilarity index. Non-metric multidimensional scaling ordination was performed using the vegan package in R (Oksanen et al. 2017). To examine the impact of stress treatment and fish size (length) on community structure, multivariate statistical analysis of community separation (PERMANOVA) was performed using Adonis in the vegan package having first checked that the data met the assumption of homogeneity of variance using the Betadisper function.

We examined the effect of stress treatment on the abundance of individual ASVs within the gut and skin microbiomes using DeSeq2 (Love et al. 2014), using rarefied data as recommended for microbiome libraries (Weiss et al. 2017). The DeSeq2 models included independent filtering of low coverage ASVs, employed default settings for outlier detection and moderation of ASV dispersion estimates, and optimised power for identification of differentially abundant ASVs at a threshold of alpha=0.05. ASVs abundance was considered significantly different at FDR <0.05.

## Results

### Microbiome alpha and beta diversity

There was no detectable effect of stress or fish size on measures on alpha diversity in the gut or skin microbiome, or in the tank water (Figure 1). For Chao1 richness there was no significant effect of stress (Gut: F_2,26_ =3.11, P=0.06; Skin: F_2,26_ =1.93, P=0.17; Water: F_2,3_=5.54, P=0.10). For Shannon diversity, there was no detectable effect of either stress treatment or fish size (P>0.4 in all cases).

**Figure 1.**
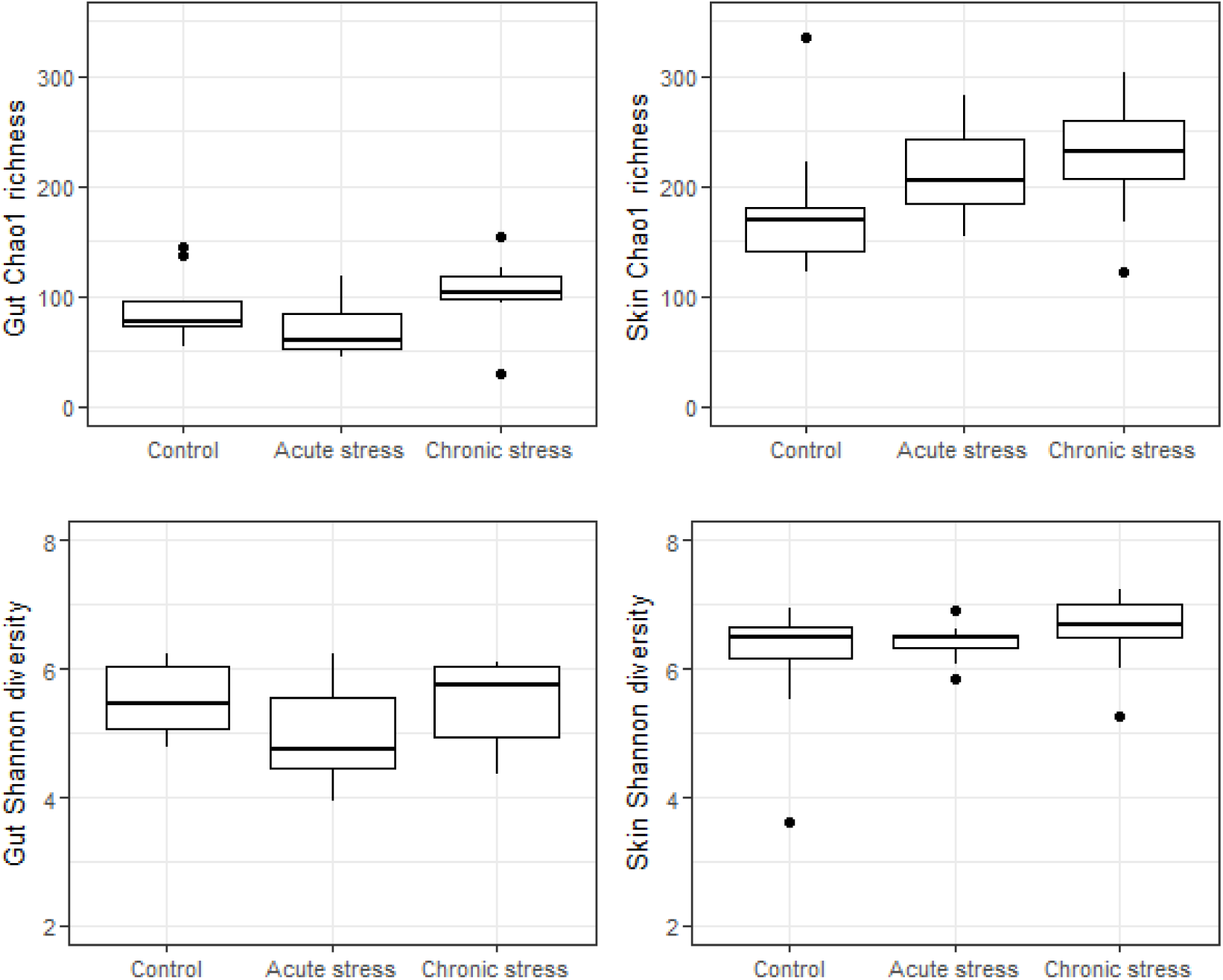
Alpha diveristy (Chao1 richness and Shannon diversity) in the gut and skin microbiome in fish exposed to an acute cold stress during embryogenesis and a chronic post-hatch environmental stress.

In contrast, we identified a pronounced effect of stress on microbiome beta diversity (Figure 2). There was a significant effect of stress, but not fish size (length), on both gut and skin community structure (Gut: *Stress F*_25,2_= 1.95, P =0.012, *Length F*_25,1_= 0.85, P =0.582; Skin: *Stress F*_25,2_=3.81, P =0.001, *Length F*_25,1_= 0.85, P =0.692). In particular, the skin microbiome of acutely stressed fish was clearly separated from that of the controls and chronically stressed groups.

**Figure 2.**
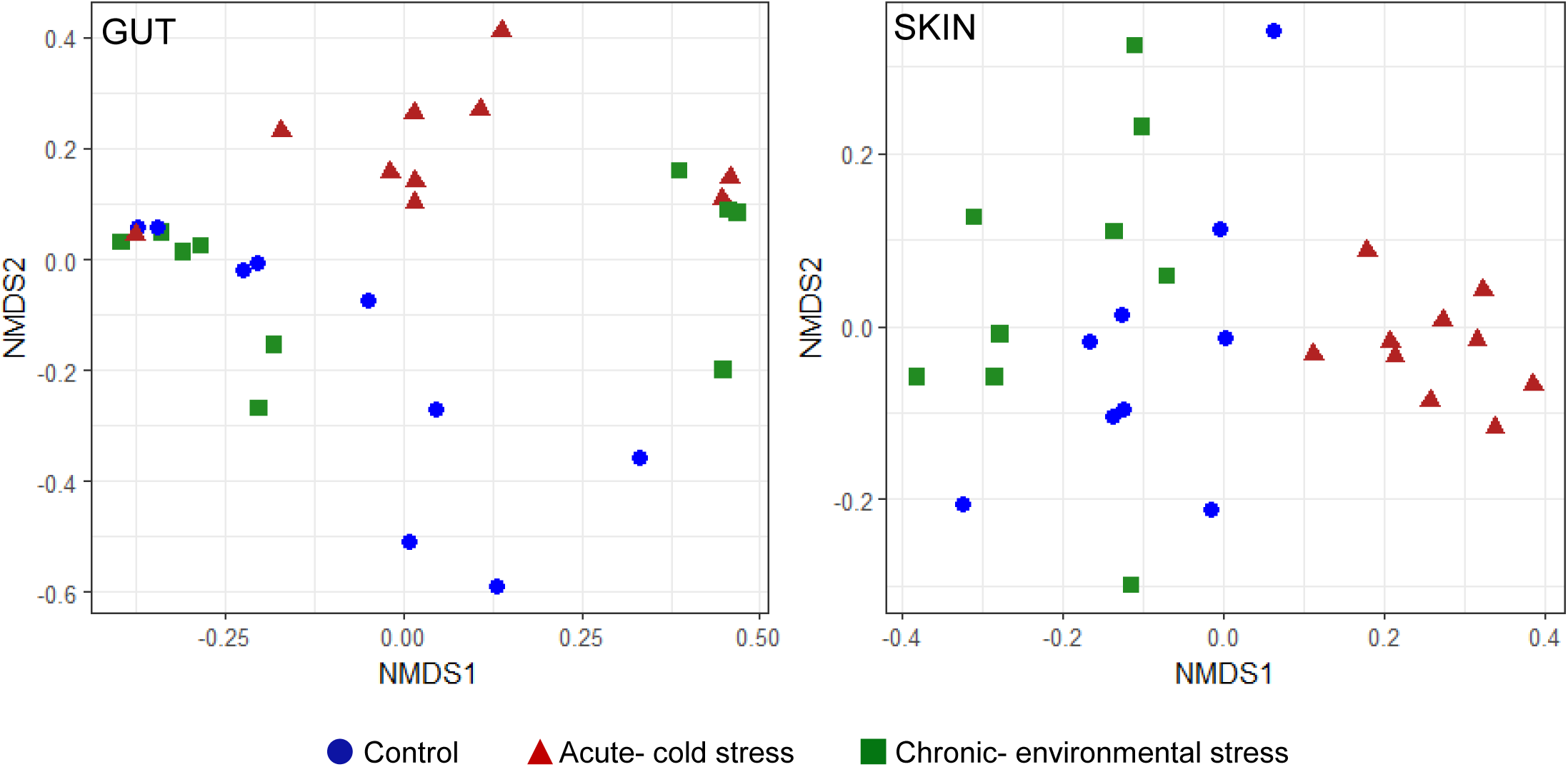
Non-metric multidimensional scaling (NMDS) ordination of microbial community structure based on Bray-Curtis distances, for all gut and skin samples.

### Microbiome composition and ASV abundance

Overall, the most abundant bacterial phyla present in the gut microbiome were Firmicutes and Proteobacteria, with lower levels of Terenicutes, Actinobacteria and Planctomycetes. The skin microbiome was dominated by Proteobacteria (mainly Gammaproteobacteria), with smaller numbers of Firmicutes, Actinobacteria and Bacteroidetes, while the tank water samples were dominated by Proteobacteria and Bacteroidetes.

There was a clear effect of both acute cold stress and chronic environmental stress on the composition of the gut microbiome. We identified 65 gut ASVs with significantly different abundance (FDR<0.05) in acutely stressed fish compared to the controls, and 32 gut ASVs that were differentially abundant between chronically-stressed and control fish. Of these, 24 (75%) were similarly affected by both types of stress (Table S1, Figure 3). Notably, 25 out of the 36 gut ASVs that were present at higher levels in acutely-stressed fish were members of the class Gammaproteobacteria and, in particular, 19 (53%) were from the genus *Acinetobacter*. Similarly, amongst the 14 gut ASVs present at higher abundance in chronically stressed fish, 11 (79%) were Gammaproteobacteria including five ASVs within the genus *Plesiomonas* and two ASVs within the genus *Acinetobacter*. Overall, *Plesiomonas* and *Acinetobacter* were amongst the most abundant gut genera in both stress groups, although there was considerable variation in their abundance between individual fish. Several *Lactobacillus* sp., *Gemmata* sp. and *Candidatus Bacilloplasma* ASVs were amongst those showing the largest decline in abundance in fish subject to both stressors, compared to the controls.

**Figure 3.**
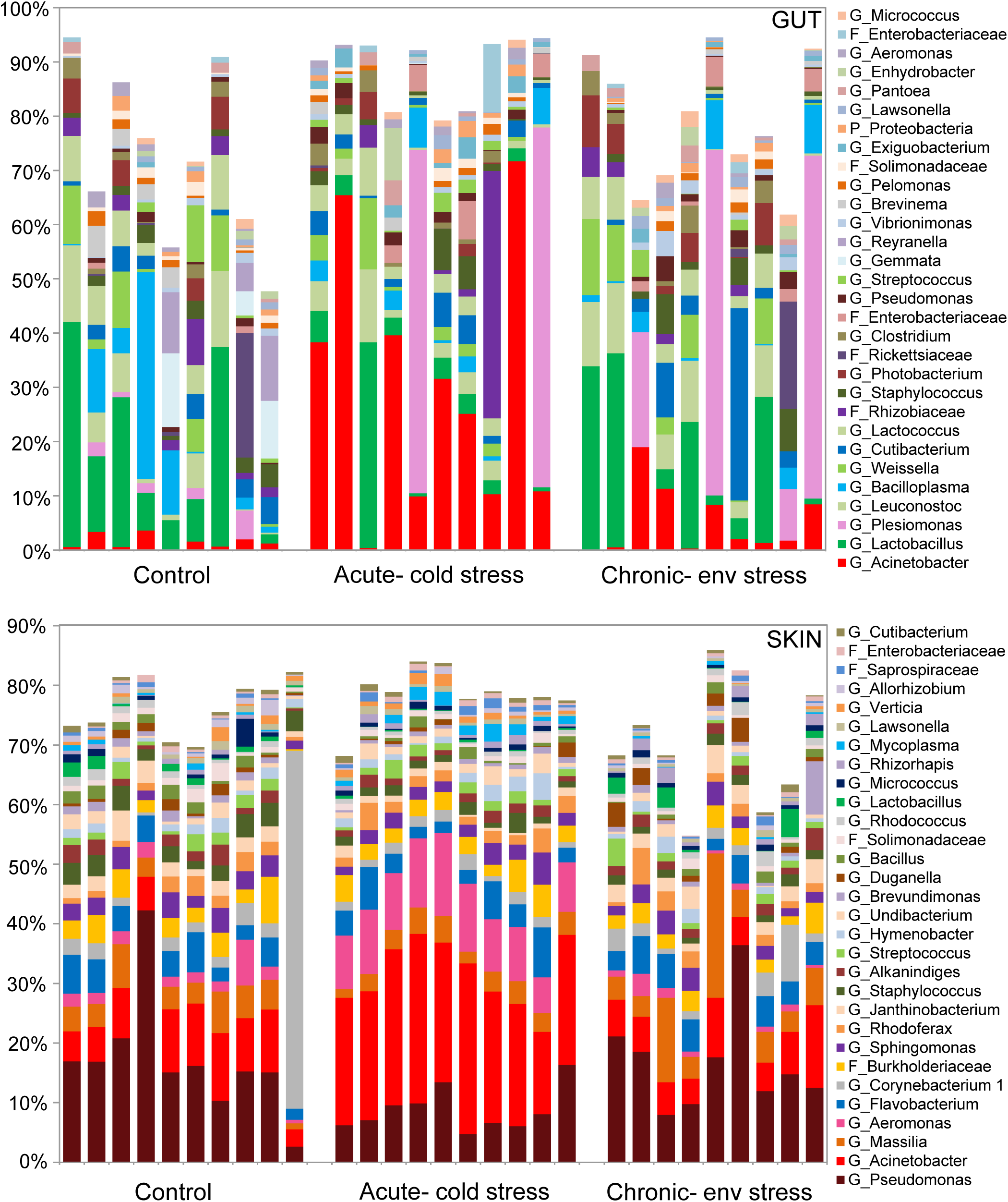
Genus-level composition of the gut and skin microbiome. Each bar represents the relative abundance of the top 30 genera, expressed as a percentage of subsampled reads.

There was also a clear effect of acute cold stress, but not of chronic stress, on the composition of the skin microbiome. We identified 87 individual skin ASVs that were present at significantly different abundance levels in acutely stressed fish compared to that of the control fish, but only one ASV was differentially abundant in fish subject to the chronic fish. Similarly to that observed in the gut, a number of Gammaproteobacteria ASVs were present at significantly higher levels in the skin of acutely stressed fish. Notably, these included four of the most abundant skin ASVs across all fish; three *Acinetobacter* sp. and one *Aeromonas* sp. (Table S2, Figure 3). However, in contrast to the gut microbiome, acute cold stress had a far more consistent effect on the skin microbiome between individual fish.

## Discussion

Our results indicate that stress experienced during early-life can have persistent effects on the diversity and structure of the Atlantic salmon microbiome, even in the absence of significant effects on survival, growth or condition factor (Uren Webster et al. 2018b). The impact of stress on the salmon microbiome was dependent on the type of stressor, as well as community type (gut or skin). However, we also identified some similarities in microbiome response to stress, suggesting there may be some common stress-signatory bacterial taxa. Acute cold stress during late embryogenesis induced very marked and consistent changes in the overall structure of the skin microbiome four months post hatch. This was characterised by the altered abundance of a large number of individual ASVs, and in particular a marked increase in several *Acinetobacter* sp. and *Aeromonas* sp., which were amongst the most abundant taxa present in the skin microbiome. In contrast, chronic, post-hatch environmental stress had few discernible effects on skin microbial community diversity or structure. The gut microbiome was altered by both stressors in a similar way, although acute cold stress had a more extensive effect. These changes were characterised by a shift from a community largely dominated by Bacilli, especially Lactobacilli, Mollicutes and Planctomycetes in the control fish, to one dominated by Gammaproteobacteria with elevated abundance of *Acinetobacter* sp. and/or *Plesiomonas* sp. in particular.

### Likely mechanisms of microbiome stress disruption

Microbiome community assembly is determined by a series of complex interactions with the host, the environment and between microbiota, which vary with age and body site (Walter & Ley 2011). Our results suggest that the mechanisms by which the acute and chronic stress affect the salmon microbiome are likely to be different, reflecting the different timing, nature and duration of each stressor, as well as the differential communities associated with the skin and the gut.

In the case of the acute stress during late embryogenesis, it is probable that exposure to iced water directly affected microbial communities associated with the egg shell, and thus the subsequent assembly of the gut and skin microbiome. The egg shell microbiome is extremely diverse and known to be directly influenced by the chemical and physical characteristics of the surrounding water, including temperature (Liu et al. 2014; Wilkins et al. 2016). Upon hatching, these microbial communities are the primary colonisers of the fish intestine and skin and therefore fundamentally influence successive community structure (Stephens et al. 2016; Wilkins et al. 2016). Differences in microbial colonisation history can alter the availability of ecological niches for successive colonisers, and can explain lasting differences in community assembly in otherwise identical environmental conditions (Fukami 2015; Sprockett et al. 2018; Walter & Ley 2011). The host-associated microbiome is extremely dynamic during these early stages of colonisation and proliferation, and is therefore likely to be particularly sensitive to environmental variation (Rea et al. 2016). Notably, both *Acinetobacter* sp. and *Aeromonas salmonicida*, are known psychrophiles (Beaz-Hidalgo & Figueras 2013; Doughari et al. 2011). Therefore it seems likely that in this case, the acute cold shock disrupted the eggshell microbiome, favouring these taxa with higher cold tolerance, which in turn altered initial colonisation of mucosal surfaces upon hatching. Once established, these taxa are then likely to have had an enhanced ability to out-compete subsequent colonisers and retain their dominant position through niche pre-emption (Sprockett et al. 2018; Walter & Ley 2011). It could be that these colonisation effects remained more pronounced and consistent in the skin microbiome because gut microbial communities are subsequently more readily influenced by the diet (Ingerslev et al. 2014; Lokesh et al. 2019).

Host stress response is likely to be an important factor underlying the effects of chronic stress on gut microbiome structure. In these same fish we also found evidence of considerable stress-related transcriptional changes, including in the expression of genes involved in glucocorticoid production and oxidative stress response, and a depressed immune response to a simulated pathogenic challenge (Uren Webster et al. 2018b). Furthermore, a similar chronic environmental stress, consisting of lack of tank enrichment, has previously been reported to cause elevated stress levels (plasma cortisol) in juvenile Atlantic salmon (Näslund et al. 2013). Therefore, elevated levels of circulating cortisol and/or stress-induced changes in the immune system may have influenced the gut microbiota. Cortisol-mediated stress response in salmonids is established at two weeks post hatching (Barry et al. 1995), and in mammals cortisol is known to directly affect microbiome diversity and composition. Specifically, elevated glucocorticoid concentrations have been shown to promote Gammaproteobacteria and inhibit probiotic taxa including Lactobacilliales (Mudd et al. 2017; Stothart et al. 2016; Zijlmans et al. 2015). This is highly consistent with the changes we observed in the gut microbiome.

### Potential implications of stress-disruption of the microbiome

While our results suggest that acute cold stress and chronic environmental stress affect the salmon microbiome via different mechanisms, both stressors favoured an increase in the abundance of certain taxa within the class Gammaproteobacteria, especially *Acinetobacter, Aeromonas* and *Plesiomonas*. The same taxa have previously been linked to hypoxia and social stress (Boutin et al. 2013; Ringø et al. 1997), suggesting they could have an inherently higher resilience to stress, and the ability to thrive in the absence of wider microbial competition (Shade et al. 2012). *Acinetobacter* and *Aeromonas* are both widely distributed in soil, water and as commensals in many animals (Doughari et al. 2011; Janda & Abbott 2010) and, crucially, also include a number of pathogenic genera that cause significant mortalities and economic loss in aquaculture (Austin & Austin 2007). Opportunistic *Aeromonas* infections are common in fish subject to stressful conditions, for example elevated temperatures, poor water quality and during spawning, typically causing septicaemia, ulceration, haemorrhages and chronic lethargy (Beaz-Hidalgo & Figueras 2013). Psychrophilic *A. salmonidicia*, usually the predominant species found in cold-water fish, is the cause of fish furunculosis which is most associated with salmonids, while mesophilic species including *A. hydrophila* and *A. veronii* also cause an assortment of stress-induced diseases in warmer-water fish (Austin & Austin 2007; Beaz-Hidalgo & Figueras 2013). Additionally, *Acinetobacter johnsonii, Acinetobacter iwoffi* and *Acinetobacter pittii* have been reported as emergent, antibiotic-resistant, opportunistic fish pathogens, with pathologies including scale loss, skin/fin erosion, haemorrhaging, intestinal inflammation and ∼20% mortality rate (Kozinska et al. 2014; Li et al. 2017; Roald & Hastein 1980).

Mucosal-associated microbial communities in the gut and on the skin provide protection against pathogens through colonisation resistance, secretion of antimicrobial compounds and stimulation of the immune system (Brestoff & Artis 2013; Kamada et al. 2013). Disrupting the integrity of this critical defence mechanism can increase risk of infection (Khosravi & Mazmanian 2013). Therefore, this increased abundance of the taxa which may include important fish pathogens in the gut following both acute and chronic stress, as well as in the skin following acute stress, suggests an enhanced risk of opportunistic infection in the case of further stress or injury. In the gut we also observed that both acute and chronic stress reduced the abundance of beneficial *Lactobacillus* sp., which has been linked to intestinal inflammation and increased susceptibility to enteric pathogens (He et al. 2013; Liu et al. 2016). Furthermore, disruption of microbiome balance through increased dominance of certain taxa, as we observed in the gut of stressed fish, is known to follow exposure to stress during early life in mammals and, critically, has also been associated with wider adverse effects on health status and immunity (Borre et al. 2014; Foster et al. 2017; Khosravi & Mazmanian 2013; Rea et al. 2016).

There were no lasting effects on survival or growth, but in parallel we also found that both of these early life stressors induced considerable transcriptional and epigenetic effects in the gills of these fish (Uren Webster et al. 2018b). Chronic stress, in particular, was associated with differential expression of a number of genes with critical immune function, as well as wider metabolic function. We also found that chronic stress impaired immune response to a subsequent pathogenic challenge while the acute stress enhanced it, and there was evidence that stress-altered mechanisms of epigenetic regulation may have contributed to these effects. Importantly this shows that both the acute and chronic stress had wider effects on the immune system. Links between intestinal microbiota and the host immune system have been well established, with the microbiome playing a critical role in the maturation and differentiation of the adaptive immune system, while immune cells also regulate microbiota assemblage (Belkaid & Hand 2014; Brown et al. 2013). While it is not possible to directly relate these effects of stress on the gut and skin microbiome with the effects on the immune system identified in the gill, our results highlight that disruption of the microbiome may potentially contribute to broader, interactive effects of stress on Atlantic salmon fry.

In summary, we found that early life stress induced persistent effects on the gut and skin microbiome of Atlantic salmon fry, in a stress, and tissue, specific manner. This highlights the importance of considering subtle, sub-lethal impacts of early-life stress on the microbiome, even in the absence of outward effects on growth and condition. While acute cold shock during late embryogenesis caused extremely pronounced changes in skin microbial community, both the acute stress and chronic post-hatch stress caused similar, but more variable, changes in the structure and diversity of the gut microbiome. Different mechanisms are likely to account for this stress and tissue specificity. Disruption of eggshell microbial communities, altering microbial community colonisation and succession is perhaps most likely to explain the effects of acute cold stress on the skin and gut microbiome, while host stress response may contribute to the effects of chronic stress in the gut. The two different stressors promoted the same ASVs within the genera *Acinetobacter* and *Aeromonas*, which include several important fish pathogens. This suggests that early life stress may increase the risk of opportunistic infections in the case of further stress or injury, which is very relevant for intensive aquaculture, where multiple stressors are commonplace (Iwama et al. 2011). Given that these taxa have also been reported to respond to other types of stress in a range of fish species (Boutin et al. 2013; Ringø et al. 1997; Zha et al. 2018), they may represent novel biomarkers of stress of potential use in aquaculture.

## Acknowledgements

We are grateful to Alastair Hamilton and Landcatch Natural Selection for supplying us with Atlantic salmon eggs, to Matthew Hitchings for conducting the Illumina sequencing and Sam Fieldwork for assistance with sampling. This work was funded by a BBSRC-NERC Aquaculture grant (BB/M026469/1) to CGL and the Welsh Government and Higher Education Funding Council for Wales (HEFCW) through the Sêr Cymru National Research Network for Low Carbon Energy and Environment (NRN-LCEE) to SC.

