## Supporting information for "Early life stress causes persistent impacts on the microbiome of Atlantic salmon"

P2-3: **Table S1**. Significantly differentially abundant ASVs in the gut.

P3-4: **Table S2.** Significantly differentially abundant ASVs in the skin.

### Table S1. Significantly differentially abundant ASVs in the gut

| ASV | Mean base abundance | Acute stress log2 fold change | Acute stress FDR | Chronic stress log2 fold change | Chronic stress FDR |
|---|---|---|---|---|---|
| G_Acinetobacter | 49.92 | -4.94 | 5.91E-12 | -1.41 | 1.18E-01 |
| G_Plesiomonas | 46.08 | -1.84 | 3.90E-02 | -1.89 | 4.50E-02 |
| G_Plesiomonas | 30.05 | -1.79 | 4.45E-02 | -2.08 | 2.52E-02 |
| G_Plesiomonas | 23.10 | -2.08 | 1.48E-02 | -2.31 | 1.09E-02 |
| G_Plesiomonas | 18.26 | -2.33 | 4.77E-03 | -2.47 | 6.03E-03 |
| G_Acinetobacter | 14.21 | -3.96 | 2.31E-09 | -1.39 | 9.01E-02 |
| G_Acinetobacter | 13.32 | -3.77 | 1.06E-08 | -1.66 | 3.38E-02 |
| G_Acinetobacter | 13.24 | -3.82 | 6.63E-09 | -1.54 | 5.13E-02 |
| G_Acinetobacter | 11.16 | -3.47 | 3.25E-07 | -1.63 | 4.40E-02 |
| G_Gemmata | 10.93 | 3.70 | 3.25E-07 | 3.70 | 1.92E-06 |
| G_ Bacilloplasma | 10.15 | 0.97 | 2.44E-01 | 2.27 | 6.03E-03 |
| G_Acinetobacter | 9.97 | -3.05 | 2.97E-05 | -1.60 | 6.34E-02 |
| G_Acinetobacter | 9.71 | -3.65 | 6.63E-09 | -1.04 | 2.48E-01 |
| G_Brevinema | 9.60 | 0.28 | 7.79E-01 | 1.88 | 2.52E-02 |
| G_Photobacterium | 9.52 | 2.16 | 5.28E-03 | 0.71 | 5.01E-01 |
| G_Exiguobacterium | 9.37 | -2.88 | 5.13E-05 | -2.06 | 1.09E-02 |
| G_Acinetobacter | 9.30 | -3.13 | 7.51E-06 | -1.60 | 5.13E-02 |
| G_Acinetobacter | 9.30 | -3.56 | 5.59E-08 | -0.70 | 4.73E-01 |
| G_Acinetobacter | 8.67 | -3.36 | 6.59E-07 | -0.87 | 3.65E-01 |
| G_Lactobacillus | 8.08 | 2.40 | 1.65E-03 | 0.04 | 9.95E-01 |
| G_Acinetobacter | 7.96 | -3.26 | 1.50E-06 | -0.78 | 4.34E-01 |
| G_Hyphomicrobium | 7.55 | 3.33 | 2.51E-06 | 3.33 | 1.21E-05 |
| G_Acinetobacter | 7.07 | -3.02 | 1.37E-05 | -0.90 | 3.55E-01 |
| F_Enterobacteriaceae | 7.03 | -2.02 | 1.28E-02 | -2.07 | 1.67E-02 |
| G_Gemmata | 6.84 | 3.23 | 4.62E-06 | 3.23 | 1.60E-05 |
| G_Acinetobacter | 6.73 | -3.09 | 2.07E-06 | -0.91 | 3.13E-01 |
| G_Acinetobacter | 6.73 | -3.20 | 1.00E-06 | -0.43 | 7.03E-01 |
| G_Acinetobacter | 6.51 | -2.99 | 7.51E-06 | -0.94 | 3.10E-01 |
| F_Enterobacteriaceae | 6.32 | -1.43 | 7.78E-02 | -2.08 | 1.12E-02 |
| G_Acinetobacter | 5.88 | -2.56 | 3.17E-04 | -1.34 | 1.28E-01 |
| G_Clostridium | 5.59 | -0.34 | 7.19E-01 | -1.69 | 3.38E-02 |
| G_Lactobacillus | 5.47 | 2.77 | 3.77E-05 | 1.55 | 3.53E-02 |
| G_Rhodococcus | 5.47 | 2.44 | 1.10E-03 | 0.96 | 3.10E-01 |
| F_Rhodobacteraceae | 5.47 | 2.97 | 2.37E-05 | 2.97 | 7.88E-05 |
| G_Brevinema | 5.36 | 2.74 | 1.10E-04 | 1.88 | 1.67E-02 |
| G_Reyranella | 5.17 | 2.91 | 3.15E-05 | 2.91 | 9.30E-05 |
| G_Exiguobacterium | 4.95 | -2.74 | 5.27E-05 | -0.30 | 7.98E-01 |
| G_Acetobacter | 4.87 | -2.33 | 1.79E-03 | -1.04 | 3.04E-01 |
| G_Reyranella | 4.72 | 2.79 | 6.69E-05 | 2.79 | 1.98E-04 |
| G_Acinetobacter | 4.43 | -2.36 | 1.16E-03 | -0.80 | 4.34E-01 |
| G_Reyranella | 4.28 | 2.67 | 1.17E-04 | 2.67 | 3.20E-04 |
| 1_Proteobacteria | 4.20 | -2.19 | 2.52E-03 | -1.06 | 2.75E-01 |
| C_Gammaproteobacteria | 4.09 | 2.62 | 1.54E-04 | 2.62 | 3.83E-04 |
| F_Holosporaceae | 3.94 | 2.58 | 1.96E-04 | 2.58 | 4.51E-04 |
| G_Pseudomonas | 3.91 | -0.98 | 2.44E-01 | -2.08 | 1.02E-02 |
| G_Tatumella | 3.83 | -2.13 | 3.76E-03 | -0.81 | 4.34E-01 |
| G_ Bacilloplasma | 3.65 | 2.08 | 4.12E-03 | 1.01 | 2.71E-01 |
| G_Acinetobacter | 3.46 | -2.04 | 4.77E-03 | -0.73 | 4.73E-01 |
| G_Plesiomonas | 3.27 | 0.02 | 9.89E-01 | -2.17 | 6.47E-03 |
| G_Flavobacterium | 3.16 | -2.17 | 2.04E-03 | 0.02 | 9.95E-01 |
| G_Fimbriiglobus | 2.97 | 2.21 | 1.33E-03 | 2.21 | 3.14E-03 |
| G_Lawsonella | 2.90 | -2.04 | 4.10E-03 | 0.02 | 9.95E-01 |
| G_Photobacterium | 2.90 | 1.50 | 3.88E-02 | 0.12 | 9.67E-01 |
| G_Vibrionimonas | 2.83 | 1.82 | 9.52E-03 | 0.96 | 2.66E-01 |
| G_Streptococcus | 2.72 | -1.80 | 1.60E-02 | -0.75 | 4.67E-01 |
| G_Pantoea | 2.49 | 0.02 | 9.89E-01 | -1.82 | 1.83E-02 |
| G_SH-PL14 | 2.49 | 1.52 | 4.30E-02 | 0.67 | 4.73E-01 |
| G_Fimbriiglobus | 2.49 | 1.96 | 3.76E-03 | 1.96 | 7.53E-03 |
| G_Paracoccus | 2.42 | -1.78 | 1.38E-02 | 0.02 | 9.95E-01 |
| G_Mycoplasma | 2.34 | 1.63 | 2.27E-02 | 1.13 | 1.65E-01 |
| G_Aeromonas | 2.34 | -1.73 | 1.55E-02 | 0.01 | 9.95E-01 |

| | | | | | |
|---|---|---|---|---|---|
| G_Paracoccus | 2.34 | -1.56 | 3.35E-02 | -0.41 | 7.07E-01 |
| G_Ulvibacter | 2.31 | 1.67 | 1.55E-02 | 1.23 | 1.10E-01 |
| G_Rothia | 2.27 | 0.01 | 9.89E-01 | -1.68 | 2.76E-02 |
| G_Pseudomonas | 2.16 | -1.60 | 2.48E-02 | 0.01 | 9.95E-01 |
| F_Pirellulaceae | 2.16 | 1.70 | 1.48E-02 | 1.70 | 2.36E-02 |
| G_Enterococcus | 2.08 | 1.64 | 1.81E-02 | 1.64 | 2.76E-02 |
| G_Mycoplasma | 2.01 | 1.54 | 3.80E-02 | 1.54 | 4.99E-02 |
| G_Streptococcus | 2.01 | -1.48 | 3.88E-02 | 0.01 | 9.95E-01 |
| G_Anoxybacillus | 1.93 | 1.54 | 2.27E-02 | 1.54 | NA |
| G_Reyranella | 1.90 | 1.44 | 4.91E-02 | 1.44 | NA |
| G_Roseococcus | 1.82 | 1.39 | 4.98E-02 | 1.39 | NA |
| G_Lactobacillus | 1.79 | 1.37 | 4.74E-02 | 1.37 | NA |

**Table S2. Significantly differentially abundant ASVs in the skin**

| ASV | Mean base abundance | Acute stress log2 fold change | Acute stress FDR | Chronic stress log2 fold change | Chronic stress FDR |
|---|---|---|---|---|---|
| G_Pseudomonas | 451.15 | 1.67 | 0.0016 | 0.37 | 0.6138 |
| G_Acinetobacter | 267.69 | -1.99 | 8.52E-11 | 0.64 | 0.4119 |
| G_Aeromonas | 227.05 | -1.86 | 0.0016 | 0.59 | 0.4318 |
| G_Acinetobacter | 220.11 | -2.26 | 1.09E-04 | 0.18 | 0.8735 |
| G_Acinetobacter | 173.39 | -1.63 | 4.49E-05 | 0.84 | 0.2449 |
| G_Janthinobacterium | 157.01 | 1.27 | 0.0015 | -0.17 | 0.7840 |
| G_Staphylococcus | 145.82 | 0.88 | 0.0120 | 0.98 | 0.2001 |
| G_Pseudorhodobacter | 22.87 | -1.99 | 0.0098 | -1.47 | 0.2976 |
| G_Mycoplasma | 18.00 | -2.99 | 3.81E-04 | 0.99 | 0.7084 |
| G_Pseudomonas | 15.48 | -4.47 | 2.14E-11 | -0.04 | 0.9921 |
| G_Acinetobacter | 14.38 | -1.89 | 0.0252 | 0.51 | 0.6729 |
| G_Massilia | 12.61 | 3.03 | 3.81E-04 | 0.33 | 0.8048 |
| G_Methylobacterium | 10.83 | 1.93 | 0.0307 | 1.06 | 0.5940 |
| G_Gemmatimonas | 10.53 | -1.78 | 0.0374 | -2.01 | 0.2245 |
| G_Massilia | 10.49 | -1.77 | 0.0374 | -2.81 | 0.3746 |
| G_Sphingobium | 10.35 | 2.96 | 3.81E-04 | 0.84 | 0.7902 |
| G_Verticia | 9.81 | -3.62 | 2.59E-06 | -0.70 | 0.5122 |
| G_Polaromonas | 9.28 | 2.36 | 0.0066 | 0.87 | 0.7911 |
| G_Paracoccus | 9.10 | -2.15 | 0.0123 | -2.16 | 0.1834 |
| F_Burkholderiaceae | 9.02 | -2.92 | 3.81E-04 | -1.97 | 0.3035 |
| F_Burkholderiaceae | 7.91 | 3.30 | 7.86E-05 | 3.27 | 0.0336 |
| G_Streptococcus | 7.58 | -2.01 | 0.0167 | 1.00 | 0.6867 |
| G_Rothia | 7.55 | -1.97 | 0.0123 | -1.74 | 0.3095 |
| G_Bacillus | 7.32 | -2.01 | 0.0233 | -1.82 | 0.2103 |
| G_Streptococcus | 7.24 | 3.24 | 7.90E-05 | 3.20 | 0.0500 |
| G_Hymenobacter | 6.91 | 2.68 | 0.0010 | 0.65 | 0.5124 |
| F_Burkholderiaceae | 6.83 | -2.27 | 0.0129 | -2.24 | 0.2144 |
| G_Allorhizobium | 6.83 | 2.64 | 0.0013 | 0.65 | 0.5217 |
| G_Pedobacter | 6.80 | 1.82 | 0.0374 | 0.32 | 0.8071 |
| G_Hyphomicrobium | 6.79 | -2.05 | 0.0159 | -2.59 | 0.1447 |
| G_Stenotrophomonas | 6.42 | 2.11 | 0.0146 | 1.16 | 0.4997 |
| G_Acinetobacter | 6.33 | 2.06 | 0.0168 | -0.31 | 0.8134 |
| G_Chryseobacterium | 6.26 | -3.10 | 7.86E-05 | -0.46 | 0.7122 |
| G_Reyranella | 6.15 | 2.32 | 0.0066 | 0.22 | 0.8905 |
| G_Virgibacillus | 5.59 | 2.70 | 0.0010 | 1.61 | 0.2554 |
| F_Burkholderiaceae | 5.51 | 2.00 | 0.0160 | 2.70 | 0.4649 |
| G_Corynebacterium | 5.47 | -1.98 | 0.0120 | -2.49 | 0.4352 |
| G_Chryseobacterium | 5.28 | -2.62 | 0.0015 | -1.05 | 0.5940 |
| G_Rickettsiella | 5.17 | -2.86 | 3.81E-04 | -0.03 | 0.9921 |
| F_Burkholderiaceae | 5.13 | 2.27 | 0.0062 | 2.70 | 0.4352 |
| F_Burkholderiaceae | 5.13 | -2.45 | 0.0034 | -1.39 | 0.6105 |
| G_Legionella | 4.98 | 2.47 | 0.0028 | 1.17 | 0.8571 |
| G_Aquabacterium | 4.95 | -2.49 | 0.0031 | -1.09 | 0.5819 |
| F_Burkholderiaceae | 4.87 | -2.77 | 6.15E-04 | -0.03 | 0.9921 |
| G_Agitococcus | 4.87 | -1.94 | 0.0227 | -2.03 | 0.2144 |

| | | | | | |
|---|---|---|---|---|---|
| G_Nitrospira | 4.42 | -2.30 | 0.0062 | -1.23 | 0.8571 |
| F_Enterobacteriaceae | 4.42 | -2.72 | 4.00E-04 | -0.02 | 0.9921 |
| G_Pseudomonas | 4.23 | -2.62 | 0.0010 | -0.02 | 0.9921 |
| F_Moraxellaceae | 4.15 | -1.69 | 0.0435 | -2.00 | 0.2144 |
| G_Neisseria | 3.81 | 2.26 | 0.0062 | 1.47 | 0.3432 |
| O_Babeliales | 3.77 | 2.01 | 0.0144 | 0.66 | 0.5052 |
| G_Pseudomonas | 3.74 | -1.84 | 0.0252 | -0.69 | 0.5122 |
| F_Burkholderiaceae | 3.70 | -2.08 | 0.0123 | -1.09 | 0.5493 |
| G_Alkanindiges | 3.58 | 2.43 | 0.0015 | 2.41 | 0.1447 |
| G_Hyphomicrobium | 3.55 | -1.98 | 0.0149 | -1.20 | 0.4286 |
| G_Pedobacter | 3.32 | 2.12 | 0.0079 | 1.43 | 0.3296 |
| G_Persicitalea | 3.24 | -2.08 | 0.0120 | -1.02 | 0.5977 |
| G_Paludibacter | 3.17 | -1.88 | 0.0167 | 0.25 | 0.8690 |
| G_Lactobacillus | 3.06 | 2.19 | 0.0055 | 2.17 | 0.1678 |
| G_Clostridium | 3.06 | 2.21 | 0.0038 | 2.20 | 0.1686 |
| G_Acinetobacter | 3.02 | -1.70 | 0.0430 | -1.11 | 0.5226 |
| G_Pseudonocardia | 2.87 | -2.09 | 0.0080 | -0.02 | 0.9921 |
| G_Hydrogenophaga | 2.87 | -1.96 | 0.0123 | -0.48 | 0.6735 |
| G_Mesorhizobium | 2.79 | 1.76 | 0.0282 | 0.93 | 0.5940 |
| F_NS11-12 | 2.76 | -2.04 | 0.0085 | -0.02 | 0.9921 |
| G_Dolosigranulum | 2.76 | -1.90 | 0.0146 | 0.08 | 0.9921 |
| O_NB1-j | 2.72 | -2.01 | 0.0107 | -0.02 | 0.9921 |
| G_Blastocatella | 2.68 | 1.65 | 0.0406 | 0.76 | 0.8171 |
| G_Schlegelella | 2.64 | 1.99 | 0.0098 | 1.97 | 0.3308 |
| F_Terrimicrobiaceae | 2.61 | 1.93 | 0.0146 | 1.91 | 0.2144 |
| G_Shewanella | 2.60 | -1.95 | 0.0120 | -0.01 | 0.9921 |
| G_Alloiococcus | 2.57 | 1.91 | 0.0149 | 1.89 | 0.2225 |
| G_Finegoldia | 2.49 | -1.67 | 0.0374 | -0.56 | 0.6107 |
| O_Nostocales | 2.45 | 1.87 | 0.0136 | 1.86 | 0.2144 |
| G_Pseudomonas | 2.42 | 1.76 | 0.0361 | 1.74 | 0.1895 |
| F_Burkholderiaceae | 2.41 | -1.83 | 0.0167 | -0.01 | 0.9921 |
| G_Nevskia | 2.34 | -1.59 | 0.0443 | -0.49 | 0.6646 |
| F_Verrucomicrobiaceae | 2.34 | -1.76 | 0.0233 | -0.01 | 0.9921 |
| G_Flavobacterium | 2.34 | 1.76 | 0.0233 | 1.75 | 0.3558 |
| G_Exiguobacterium | 2.30 | 1.74 | 0.0252 | 1.73 | 0.3808 |
| G_Arenimonas | 2.19 | -1.66 | 0.0332 | -0.01 | 0.9921 |
| G_Actinobacillus | 2.19 | -1.66 | 0.0332 | -0.01 | 0.9921 |
| G_Chryseobacterium | 2.15 | 1.63 | 0.0361 | 1.62 | 0.1895 |
| F_Burkholderiaceae | 2.15 | -1.77 | 0.0233 | -0.12 | 0.9573 |
| G_AAP99 | 2.08 | 1.57 | 0.0423 | 1.56 | 0.2157 |
| G_Cereibacter | 2.08 | 1.53 | 0.0496 | 1.40 | 0.3296 |
| G_Perlucidibaca | 2.04 | -1.55 | 0.0406 | -0.01 | 0.9921 |
